## Supplementary material for "Independent regulation of Z-lines and M-lines during sarcomere assembly in cardiac myocytes revealed by the automatic image analysis software sarcApp": Figure Supplement with Legends

A yoU-Net Architecture

| | |
|---|---|
| Feature Maps | 16 (αa2, myom)/32 (titin) |
| Loss Function | BCE with Logits Loss |
| Feature Map Increase Factor | 3 |
| Downsampling Factor | [(2,2),(2,2),(2,2),(2,2)] |
| Activation | ReLU |
| Constant Upsampling | True |
| Optimizer | Adam |
| Steps per Epoch | 10 |
| Batch Size | 10 |

B Dice Coefficient Formula

$$\frac{2^*(\text{Prediction} \cap \text{Target})}{\sum \text{Prediction} + \sum \text{Target}}$$

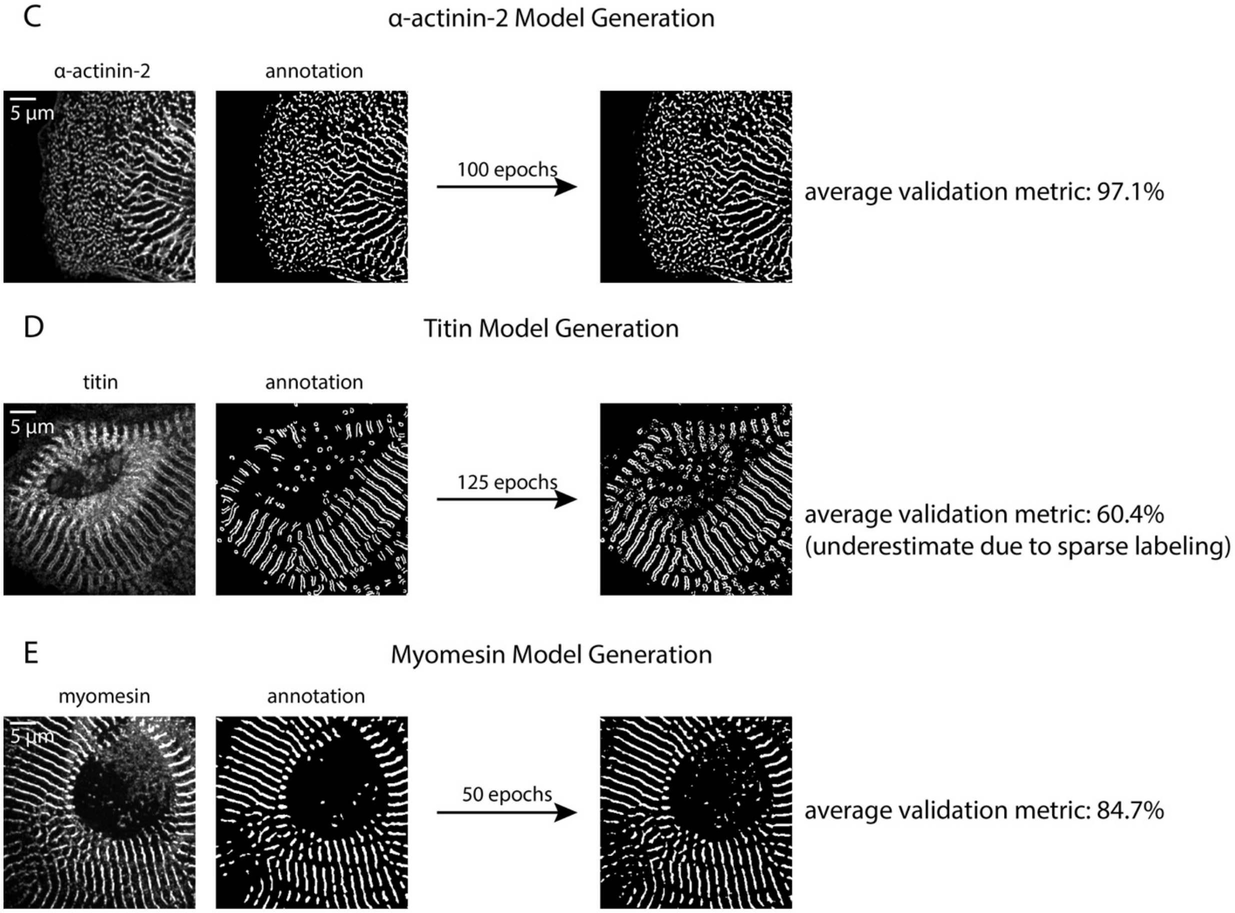


**Figure 1-figure supplement 1: Details of yoU-Net architecture and trained model generation**

A) U-Net architecture details used for this manuscript (details can be adjusted in yoU-Net). B) Dice coefficient formula, used for validating trained models. C) Representative fluorescence image and matched annotated binary of an hiCM with α-actinin-2 localized. D) Representative fluorescence image and matched annotated binary of an hiCM with titin localized. E) Representative fluorescence image and matched annotated binary of an hiCM with myomesin localized.

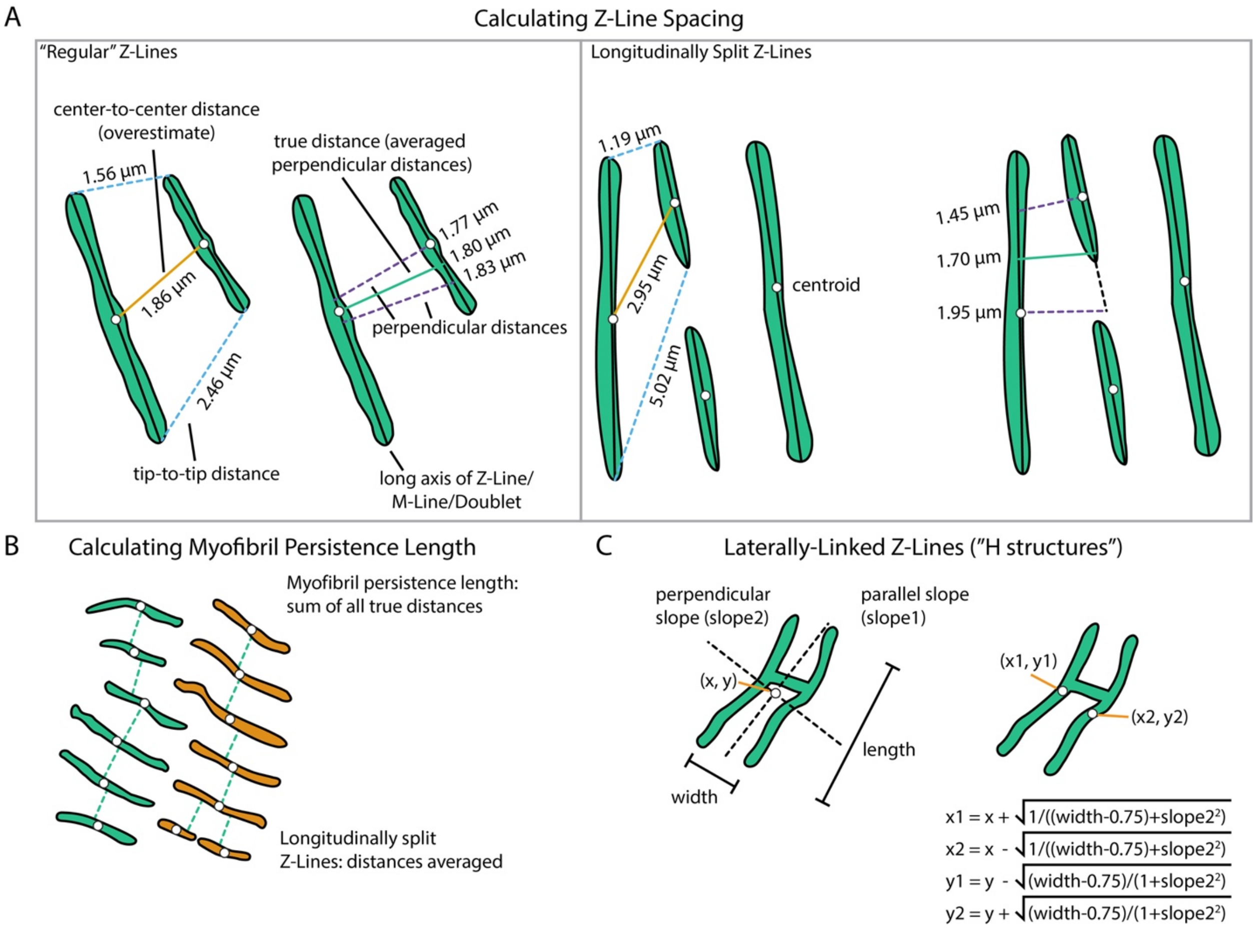


**Figure 2-figure supplement 1: Details of geometric calculations used in sarcApp**

A) How Z-Line spacing is calculated in sarcApp. In short, for each pair of Z-Lines, a line is measured from the centroid of one Z-Line to the next Z-Line using a line drawn perpendicularly to the first Z-Line. The same is done vice versa: a line from the second Z-Line to the first is measured. The averaged perpendicular distances comprise the "true" distance. B) How myofibril persistence length is calculated. In short, persistence length is the sum of all true Z-Line distances across an entire myofibril. If Z-Lines are longitudinally split, the Z-Line distances are averaged. C) How laterally-linked Z-Lines (H structures) are solved in sarcApp. In short, the length, width, and centroid of the H structure are used to back calculate the predicted centroid of each individual Z-Line.

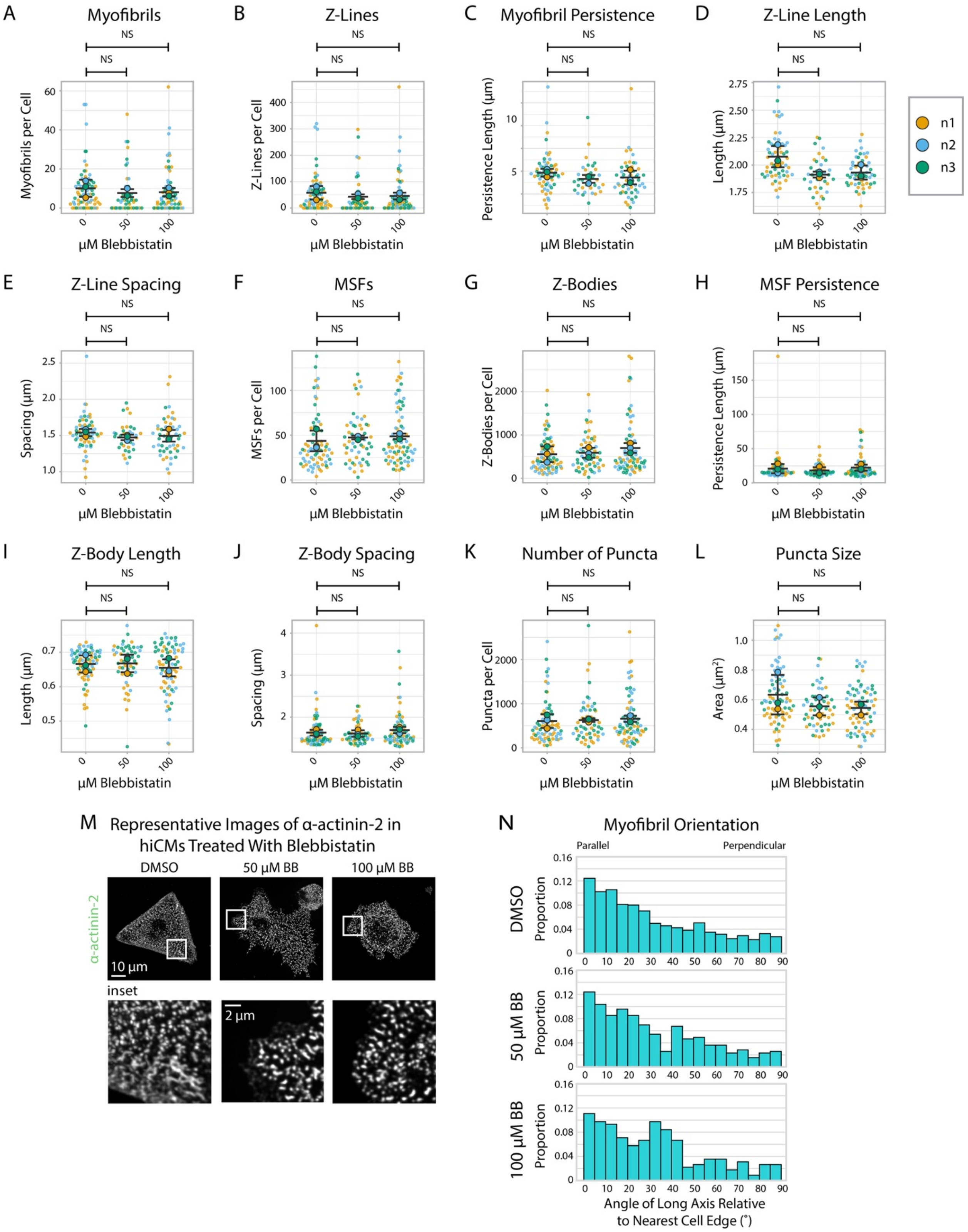


**Figure 3-figure supplement 1: α-actinin-2 quantification and organization in Blebbistatin-treated hiCMs 6 hours post-plating**

A) Myofibrils per cell in cells treated with DMSO or 50/100 μM Blebbistatin. N=3 biological replicates, 73 control cells, 54 50 μM Blebbistatin cells, and 72 100 μM Blebbistatin cells. Myofibrils defined as having 4 or more Z-Lines in a row. Details in Figure 2-figure supplement 1. B) Z-Lines per cell in hiCMs from (A). C) Average myofibril persistence length per cell in hiCMs from (B). N=59 control cells, 34 50 μM Blebbistatin cells, and 51 100 μM Blebbistatin cells (only cells with myofibrils were quantified for (C-E). D) Average Z-Line length per cell in hiCMs from (C). E) Average spacing between Z-Lines per cell in hiCMs from (C). F) MSFs per cell in hiCMs from (A). G) Z-Bodies per cell in hiCMs from (A). H) Average MSF persistence length per cell in hiCMs from (A). N=3 biological replicates, 73 control cells, 54 50 μM Blebbistatin cells, and 72 100 μM Blebbistatin cells (only cells with MSFs were quantified for (H-J). I) Average Z-Body length per cell in hiCMs from (H). J) Average spacing between Z-Bodies per cell in hiCMs from (H). K) Number of α-actinin-2-positive puncta per cell in hiCMs from (A). L) Average size of α-actinin-2-positive puncta in hiCMs from (A). M) Representative images of α-actinin-2 and F-actin in control and Blebbistatin-treated hiCMs. N) Myofibril orientation relative to the cell edge segment closest to the myofibril center, perpendicularly. N=3 biological replicates, 668 control myofibrils, 385 50 μM Blebbistatin myofibrils, and 582 100 μM Blebbistatin myofibrils. Details of quantification found in Figure 2M-O.

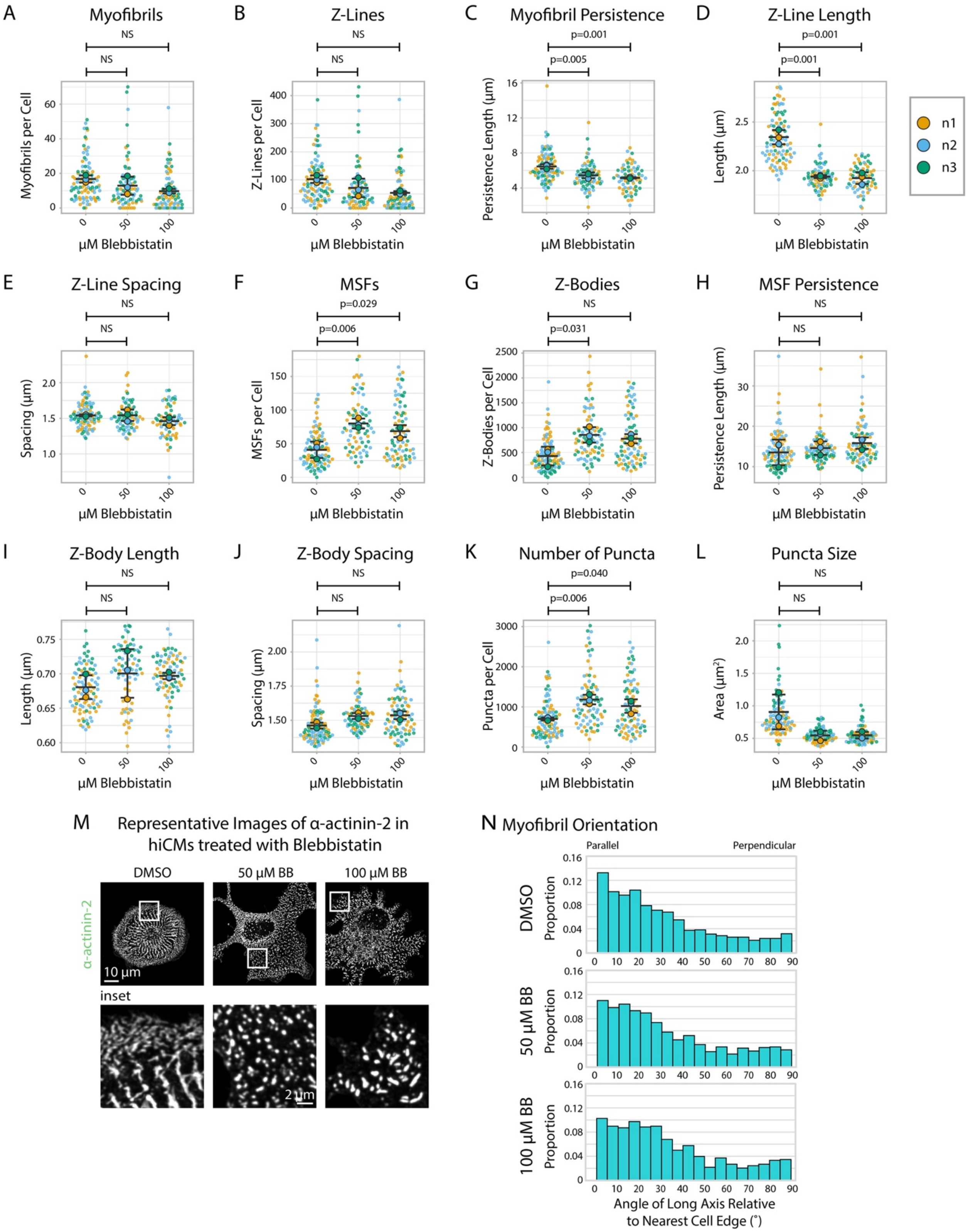


**Figure 3-figure supplement 2: α-actinin-2 quantification and organization in Blebbistatin-treated hiCMs 12 hours post-plating**

A) Myofibrils per cell in cells treated with DMSO or 50/100 μM Blebbistatin. N=3 biological replicates, 95 control cells, 77 50 μM Blebbistatin cells, and 80 100 μM Blebbistatin cells. Myofibrils defined as having 4 or more Z-Lines in a row. More quantification details found in Figure 2-figure supplement 1. B) Z-Lines per cell in hiCMs from (A). C) Average myofibril persistence length per cell in hiCMs from (A): N=3 biological replicates, 92 control cells, 69 50 μM Blebbistatin cells, and 65 100 μM Blebbistatin cells (only cells with myofibrils were quantified for (C-E). D) Average Z-Line length per cell in hiCMs from (C). E) Average spacing between Z-Lines per cell in hiCMs from (C). F) MSFs per cell in hiCMs from (A). G) Z-Bodies per cell in hiCMs from (A). H) Average MSF persistence length per cell in hiCMs from (A): N=3 biological replicates, 94 control cells, 77 50 μM Blebbistatin cells, and 80 100 μM Blebbistatin cells (only cells with MSFs were quantified for (H-J). I) Average Z-Body length per cell in hiCMs from (H). J) Average spacing between Z-Bodies per cell in hiCMs from (H). K) Number of α-actinin-2-positive puncta per cell in hiCMs from (A). L) Average size of α-actinin-2-positive puncta in hiCMs from (A). M) Representative images of α-actinin-2 and F-actin in control and Blebbistatin-treated hiCMs. N) Myofibril orientation relative to the cell edge segment closest to the myofibril center, perpendicularly. N=3 biological replicates, 1560 control myofibrils, 1012 50 μM Blebbistatin myofibrils, and 777 100 μM Blebbistatin myofibrils. Details of quantification found in Figure 2M-O.

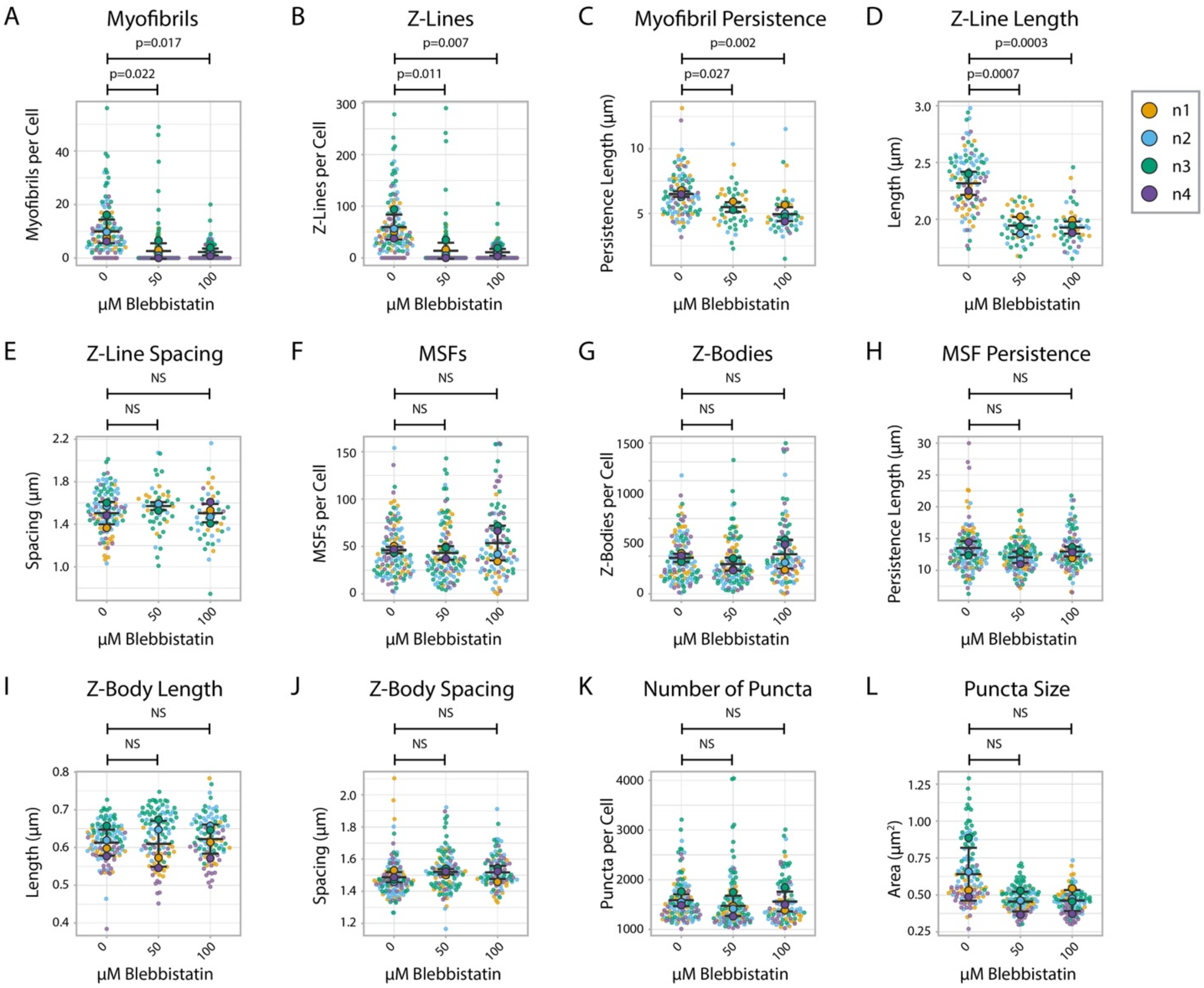


**Figure 3-figure supplement 3: α-actinin-2 quantification and organization in Blebbistatin-treated hiCMs 24 hours post-plating**

A) Myofibrils per cell in cells treated with DMSO or 50/100 µM Blebbistatin. N=4 biological replicates, 118 control cells, 108 50 µM Blebbistatin cells, and 93 100 µM Blebbistatin cells. Note that some of these subfigures were used also in Figure 3. Myofibrils defined as having 4 or more Z-Lines in a row. More quantification details found in Figure 2-figure supplement 1. B) Z-Lines per cell in hiCMs from (A). C) Average myofibril persistence length per cell in hiCMs from (A). D) Average Z-Line length per cell in hiCMs from (B): N=4 biological replicates, 104 control cells, 45 50 µM Blebbistatin cells, and 45 100 µM Blebbistatin cells (only cells with myofibrils were quantified for C-E). E) Average spacing between Z-Lines per cell in hiCMs from (C). F) MSFs per cell in hiCMs from (A). G) Z-Bodies per cell in hiCMs from (A). H) Average MSF persistence length per cell in hiCMs from (A): N=4 biological replicates, 118 control cells, 108 50 µM Blebbistatin cells, and 93 100 µM Blebbistatin cells (only cells with MSFs were quantified for (H-J). I) Average Z-Body length per cell in hiCMs from (H). J) Average spacing between Z-Bodies per cell in hiCMs from (H). K) Number of α-actinin-2-positive puncta per cell in hiCMs from (A). L) Average size of α-actinin-2-positive puncta in hiCMs from (A).

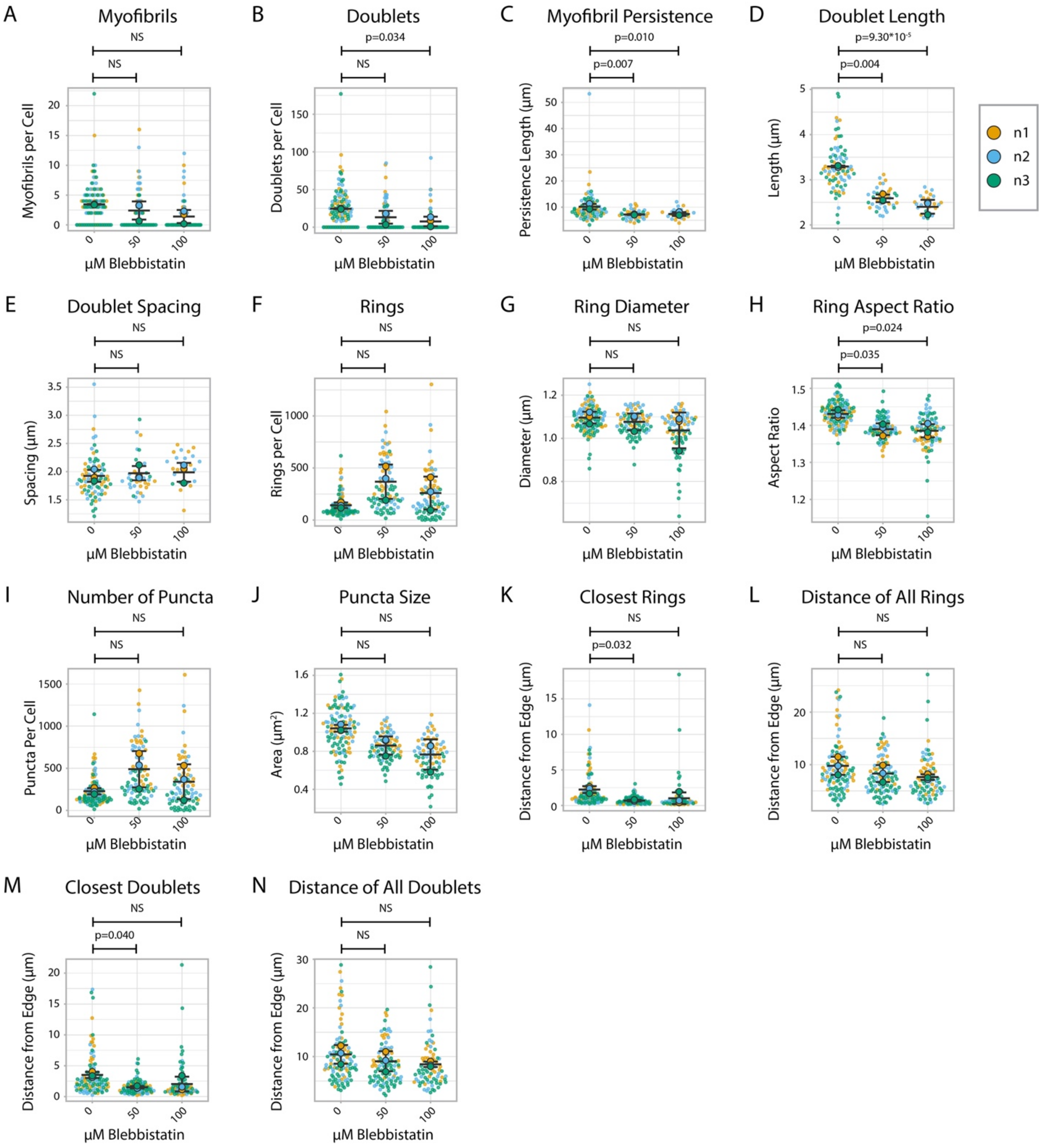


**Figure 5-figure supplement 1: Titin quantification and organization in Blebbistatin-treated hiCMs 24 hours post-plating**

A) Myofibrils per cell in cells treated with DMSO or 50/100 µM Blebbistatin. Myofibrils defined as having 4 or more doublets in a row. N=3 biological replicates, 107 DMSO control cells, 95 50 µM Blebbistatin cells, and 84 100 µM Blebbistatin cells. B) Titin doublets per cell. Only doublets within myofibrils are counted. C) Myofibril persistence length average per cell in hiCMs from S6A: N=3 biological replicates, 58 control cells, 32 50 µM Blebbistatin cells, and 21 100 µM Blebbistatin cells (only cells with myofibrils were quantified for C-E and M-N). D) Average doublet length per cell in hiCMs from (C). E) Average spacing between doublets per cell in hiCMs from (C). F) Number of titin precursor rings per cell in hiCMs from (A). G) Average ring diameter per cell per cell in hiCMs from S6A: N=3 biological replicates, 105 control cells, 95 50 µM Blebbistatin cells, and 78 100 µM Blebbistatin cells (only cells with rings were quantified for (G-H and K-L). H) Average ring aspect ratio per cell in hiCMs from (G). An aspect ratio of 1 is circular, and higher ratios are elongated. I) Total number of titin-positive puncta per cell in hiCMs from (G). J) Average size of all titin-positive puncta per cell in hiCMs from (A). K) Average distance from the edge of the five closest titin rings, per cell, in cells from (H). L) Average distance from the edge of all titin rings per cell, in cells from (H). M) Average distance from the edge of the five closest titin doublets, per cell, in hiCMs from (C). N) Average distance from the edge of all titin doublets within myofibrils per cell, in hiCMs from (C).

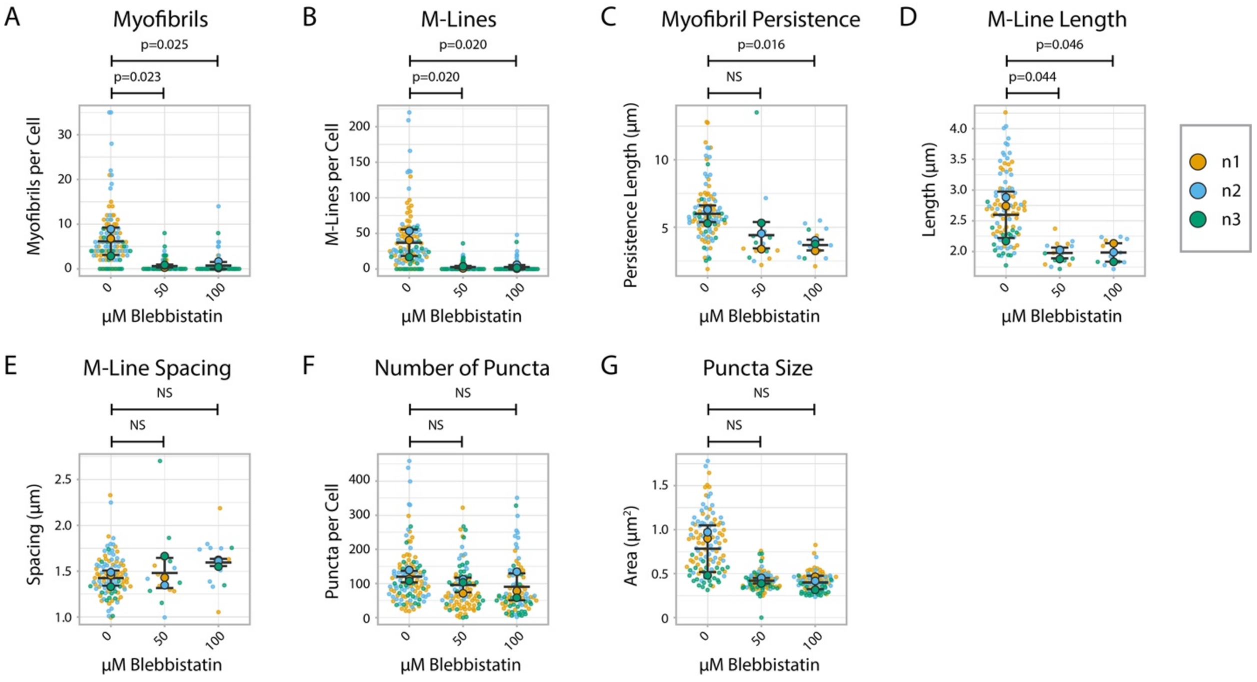


**Figure 6-figure supplement 1: Myomesin quantification and organization in Blebbistatin-treated hiCMs 24 hours post-plating**

A) Myofibrils per cell in cells treated with DMSO or 50/100 µM Blebbistatin. Myofibrils defined as having 3 or more M-Lines in a row. N=3 biological replicates, 112 DMSO control cells, 90 50 µM Blebbistatin cells, and 89 100 µM Blebbistatin cells. Quantification scheme details in Figure S2. B) M-Lines per cell from (A). C) Average myofibril persistence length per cell in hiCMs from (A): N=3 biological replicates, 97 DMSO control cells, 16 50 µM Blebbistatin cells, and 13 100 µM Blebbistatin cells. (Only cells with myofibrils were quantified for C-E). D) Average M-Line length per cell in hiCMs from (C). E) Average spacing between M-Lines per cell in hiCMs from (C). F) Number of myomesin-positive puncta per cell in hiCMs from (A). G) Average size of all myomesin-positive puncta per cell in hiCMs from (A).

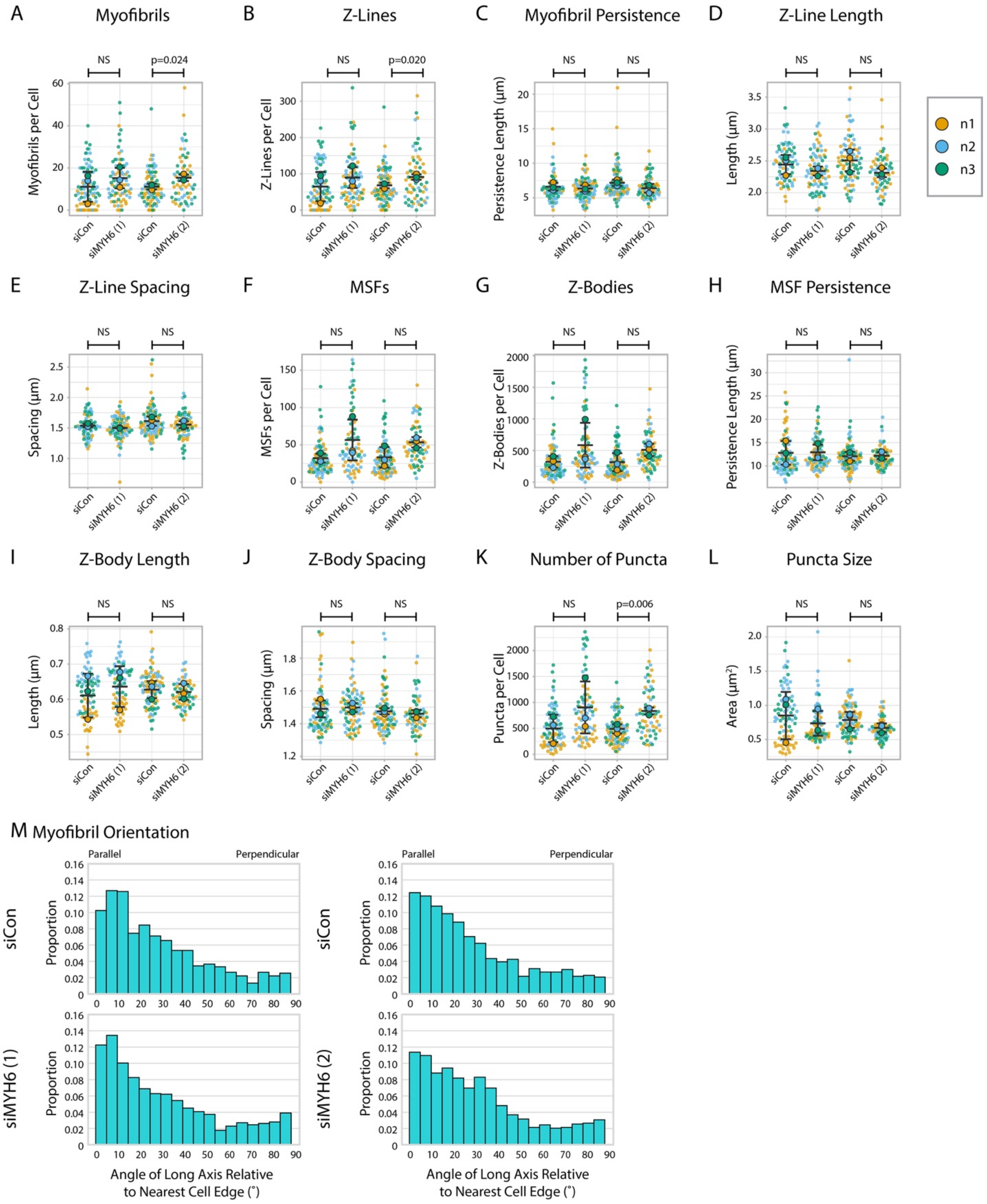


**Figure 7-figure supplement 1: α-actinin-2 quantification and organization in MYH6 knockdown hiCMs**

A) Myofibrils per cell in two groups of siCon (scramble)-treated hiCMs and two separate MYH6 siRNA-treated hiCMs (sequences 1 and 2). N=3 biological replicates, 81 siCon cells and 78 siMYH6 (1) cells, and 88 siCon cells and 63 siMYH6 (2) cells. Myofibrils defined as having 4 or more Z-Lines in a row. Quantification details found in Figure 2-figure supplement 1. B) Z-Lines per cell in hiCMs from (A). C) Average myofibril persistence length per cell in hiCMs from (B): N=68 siCon cells and 75 siMYH6 (1) cells, and 83 siCon cells and 62 siMYH6 (2) cells (only cells with myofibrils were quantified for C-E). D) Average Z-Line length per cell in hiCMs from (C). E) Average spacing between Z-Lines per cell in hiCMs from (C). F) MSFs per cell in cells from (A). G) Z-Bodies per cell in hiCMs from (A) . H) Average MSF persistence length per cell in hiCMs from (A): N=3 biological replicates, X7 siCon cells, X8 siMYH6 (1) cells, and X9 siMYH6 (2) cells (only cells with MSFs were quantified for H-J). I) Average Z-Body length per cell in hiCMs from (H). J) Average spacing between Z-Bodies per cell in hiCMs from (H). K) Number of α-actinin-2-positive puncta per cell in hiCMs from (A). L) Average size of α-actinin-2-positive puncta per cell in hiCMs from (A). M) Myofibril orientation relative to the cell edge segment closest to the myofibril center, perpendicularly. N=3 biological replicates: 898 siCon myofibrils and 1173 siMYH6 (1) myofibrils, and 964 siCon myofibrils and 974 siMYH6 (2) myofibrils. Details of quantification found in Figure 2M-O. N=3 biological replicates.

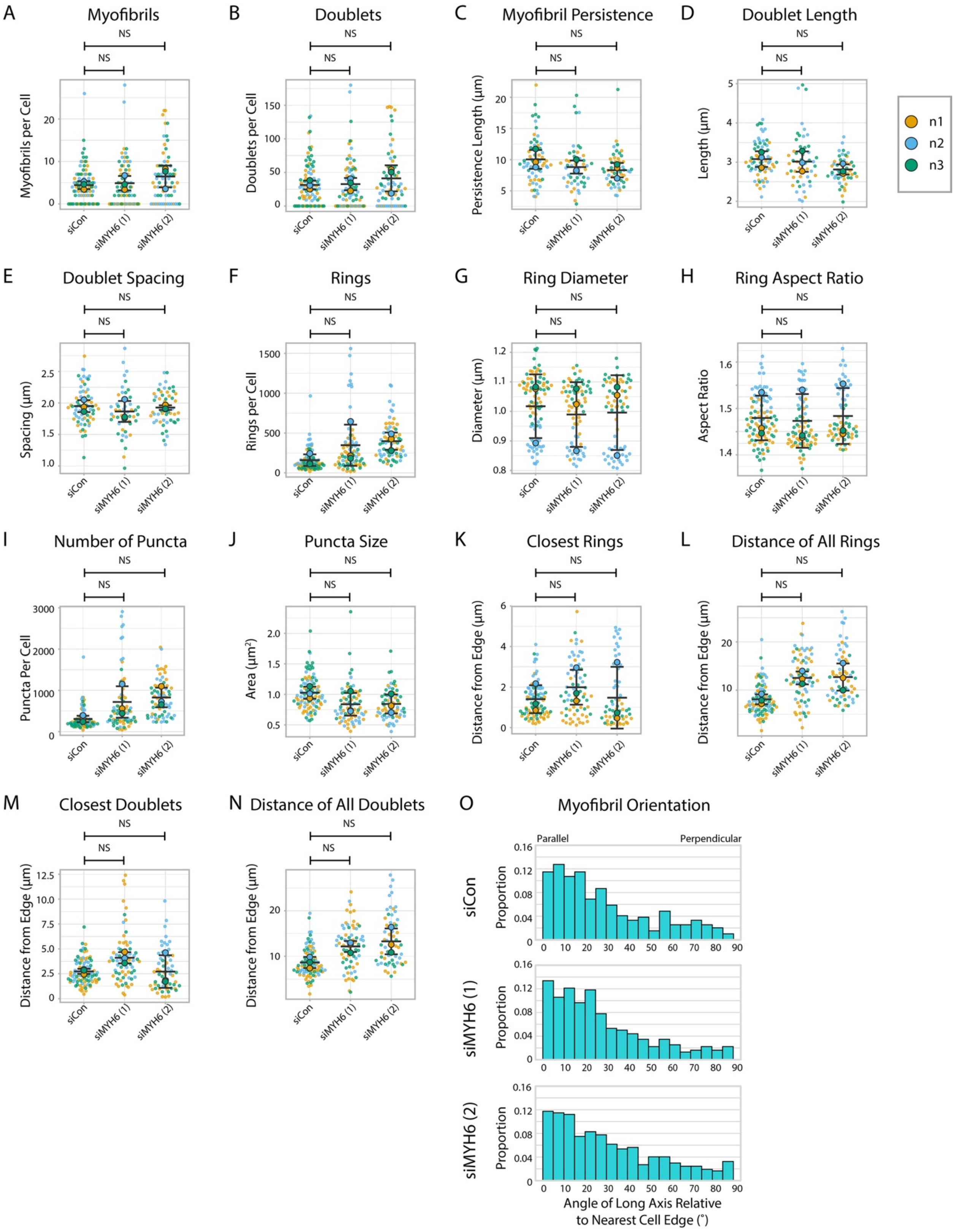


**Figure 7-figure supplement 2: Titin quantification and organization in MYH6 knockdown hiCMs**

A) Myofibrils per cell in siCon (scramble)-treated hiCMs and two separate MYH6 siRNA-treated hiCMs (sequences 1 and 2). Myofibrils were defined as having 4 or more doublets in a row. N=3 biological replicates, 91 siCon cells, 73 siMYH6 (1) cells, and 66 siMYH6 (2) cells. B) Titin doublets per cell from Figure S9A. C) Myofibril persistence length average per cell in hiCMs from (A): N=3 biological replicates, 71 siCon cells, 49 siMYH6 (1) cells, and 50 siMYH6 (2) cells (only cells with myofibrils were quantified for C-E and M, N). D) Average doublet length per cell in hiCMs from (C). E) Average spacing between doublets per cell in hiCMs from (C). F) Number of titin precursor rings per cell in hiCMs from (A). G) Average ring diameter per cel in hiCMs from (A). N=3 biological replicates, 78 siCon cells, 67 siMYH6 (1) cells, 60 siMYH6 (2) cells. (Only cells with rings were quantified for G-H and K, L). H) Average ring aspect ratio per cell in hiCMs from (G). An aspect ratio of 1 is circular, and higher ratios are elongated. I) Total number of titin-positive puncta per cell in hiCMs from (A). J) Average size of all titin-positive puncta per cell in hiCMs from (A). K) Average distance from the edge of the five closest titin rings, per cell, in hiCMs from (G). L) Average distance from the edge of all titin rings per cell in hiCMs from (G). M) Average distance from the edge of the five closest titin doublets, per cell, in hiCMs from (C). N) Average distance from the edge of all titin doublets within myofibrils per cell in hiCMs from (C). O) Myofibril orientation relative to the cell edge segment closest to the myofibril center, perpendicularly in hiCMs from (B). N=3 biological replicates, 449 siCon myofibrils, 440 siMYH6 (1) myofibrils, and 442 siMYH6 (2) myofibrils. Details of quantification found in Figure 2M-O.

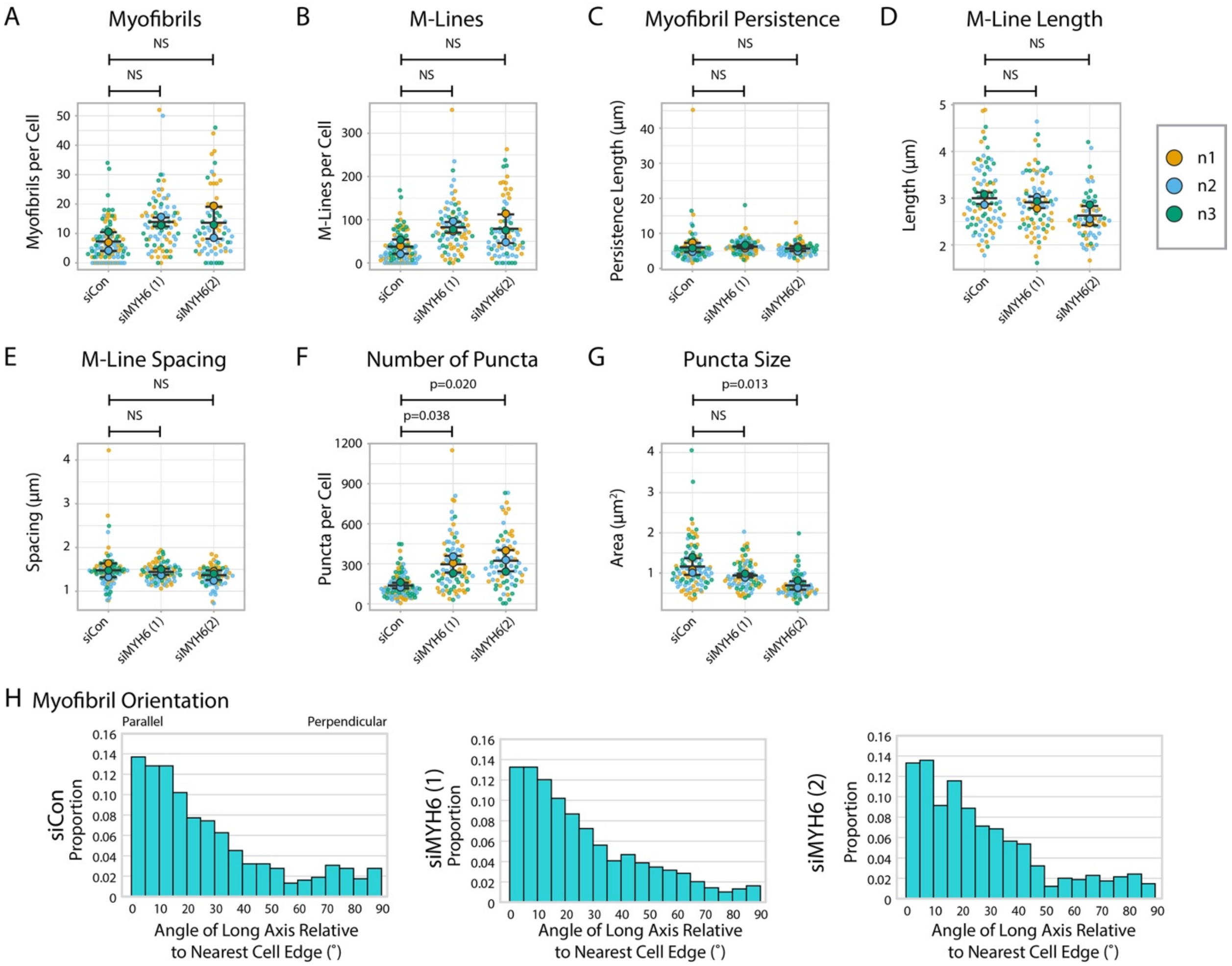


### Figure 7-figure supplement 3: Myomesin quantification and organization in MYH6 knockdown cells

A) Myofibrils per cell in siCon (scramble)-treated hiCMs and two separate MYH6 siRNA-treated hiCMs (sequences 1 and 2). Myofibrils defined as having 3 or more M-Lines in a row. N=3 biological replicates, 100 siCon cells, 80 siMYH6 (1) cells, and 71 siMYH6 (2) cells. Quantification scheme details in Figure 2-figure supplement 1. B) M-Lines per cell from (A). C) Average myofibril persistence length per cell in hiCMs from (A): N=3 biological replicates, 85 siCon cells, 78 siMYH6 (1) cells, and 64 siMYH6 (2) cells. (Only cells with myofibrils were quantified for C-E). D) Average M-Line length per cell in hiCMs from (C). E) Average spacing between M-Lines per cell in hiCMs from (C). F) Number of myomesin-positive puncta per cell in hiCMs from (A). G) Average size of all myomesin-positive puncta per cell in hiCMs from (A). H) Myofibril orientation relative to the cell edge segment closest to the myofibril center, perpendicularly. N=3 biological replicates, 685 siCon myofibrils, 979 siMYH6 (1) myofibrils, and 743 siMYH6 (2) myofibrils. Details of quantification found in Figure 2M-O.

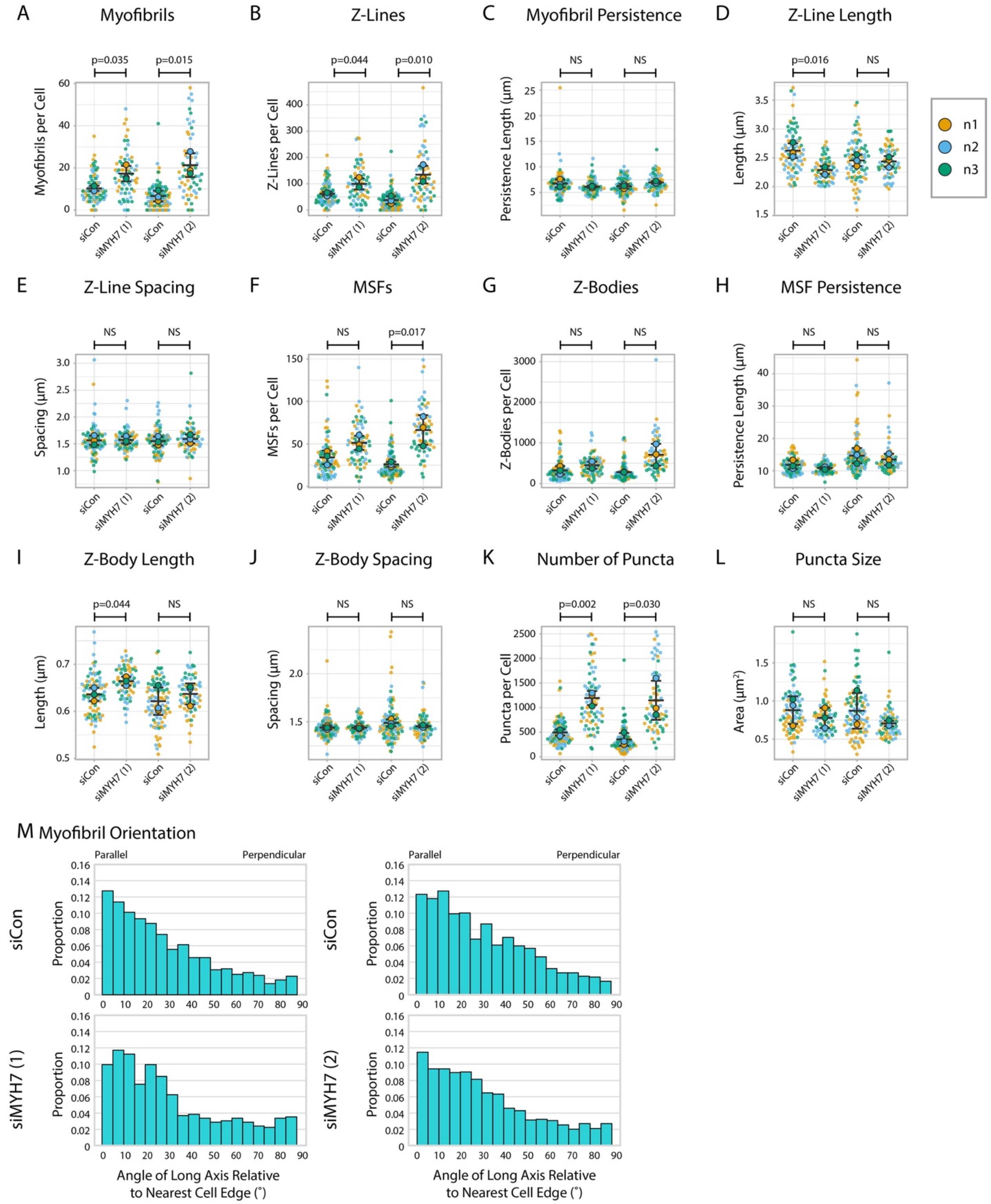


**Figure 7-figure supplement 4: α-actinin-2 quantification and organization in MYH7 knockdown hiCMs**

A) Myofibrils per cell in two groups of siCon (scramble)-treated hiCMs and two separate MYH7 siRNA-treated hiCMs (pools 1 and 2). N=3 biological replicates, 86 siCon cells and 66 siMYH7

(1) cells, and 97 siCon cells and 62 siMYH7 (2) cells. Myofibrils were defined as having 4 or more Z-Lines in a row. Details can be found in Figure 2-figure supplement 1. B) Z-Lines per cell in hiCMs from (A). C) Average myofibril persistence length per cell in hiCMs from (A): N=3 biological replicates, 81 siCon cells and 62 siMYH7 (1) cells, and 81 siCon cells and 59 siMYH7 (2) cells. (Only cells with myofibrils were quantified for C-E). D) Average Z-Line length per cell in hiCMs from (C). E) Average spacing between Z-Lines per cell in hiCMs from (C). F) MSFs per cell in cells from (A). G) Z-Bodies per cell in hiCMs from (A). H) Average MSF persistence length per cell in hiCMs from (A): N=3 biological replicates, 86 siCon cells and 66 siMYH7 (1) cells, and 97 siCon cells and 62 siMYH7 (2) cells. (Only cells with MSFs were quantified for H-J). I) Average Z-Body length per cell in hiCMs from (H). J) Average spacing between Z-Bodies per cell in hiCMs from (H). K) Number of α-actinin-2-positive puncta per cell in hiCMs from (A). L) Average size of α-actinin-2-positive puncta per cell in hiCMs from (A). M) Myofibril orientation relative to the cell edge segment closest to the myofibril center, perpendicularly. N=3 biological replicates, 878 siCon myofibrils and 1126 siMYH7 (1) myofibrils, and 622 siCon myofibrils and 1324 siMYH7 (2) myofibrils. Details of quantification found in Figure 2M-O.

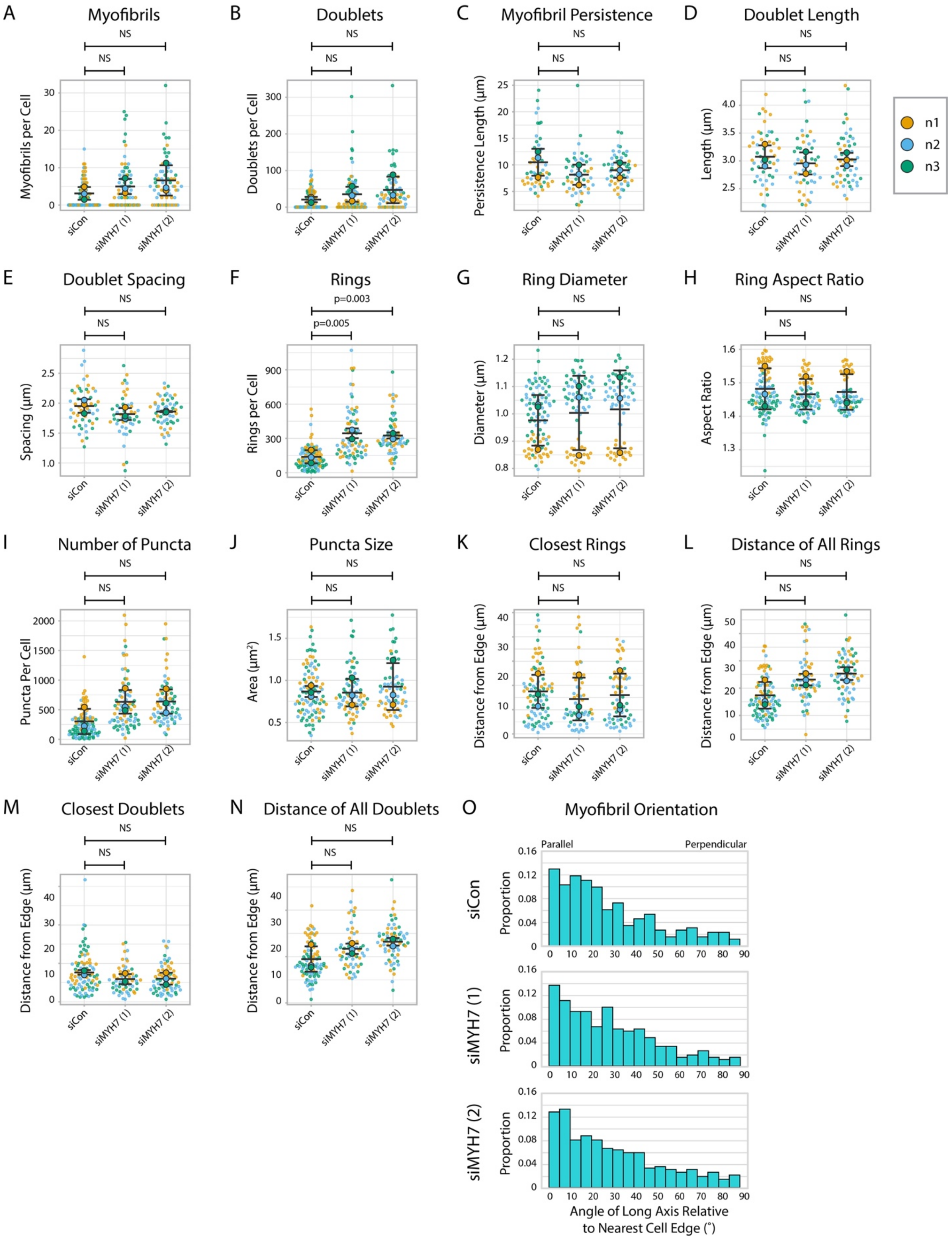


**Figure 7-figure supplement 5: Titin quantification and organization in MYH7 knockdown hiCMs**

A) Myofibrils per cell in siCon (scramble)-treated hiCMs and two separate MYH7 siRNA-treated hiCMs (pools 1 and 2). Myofibrils defined as having 4 or more doublets in a row. N=3 biological replicates, 94 siCon cells, 72 siMYH7 (1) cells, and 66 siMYH7 (2) cells. B) Titin doublets per cell from (A). C) Myofibril persistence length average per cell in hiCMs from (A): N=3 biological replicates, 56 siCon cells, 56 siMYH7 (1) cells, and 52 siMYH7 (2) cells. (Only cells with myofibrils were quantified for Figure C-E and M, N). D) Average doublet length per cell in hiCMs from (C). E) Average spacing between doublets per cell in hiCMs from (C). F) Number of titin precursor rings per cell in hiCMs from (A). G) Average ring diameter per cell in hiCMs from (A): N=3 biological replicates, 90 siCon cells, 58 siMYH7 (1) cells, and 64 siMYH7 (2) cells. (Only cells with rings were quantified for G-H and J, K). H) Average ring aspect ratio per cell in hiCMs from (G). An aspect ratio of 1 is circular, and higher ratios are elongated. I) Total number of titin-positive puncta per cell in hiCMs from (A). J) Average size of all titin-positive puncta per cell in hiCMs from (A). K) Average distance from the edge of the five closest titin rings, per cell, in hiCMs from (G). L) Average distance from the edge of all titin rings per cell in hiCMs from (G). M) Average distance from the edge of the five closest titin doublets, per cell, in hiCMs from (C). N) Average distance from the edge of all titin doublets within myofibrils per cell in hiCMs from (C). O) Myofibril orientation relative to the cell edge segment closest to the myofibril center, perpendicularly in hiCMs from (B). N=3 biological replicates, 278 siCon myofibrils, 318 siMYH7 (1) myofibrils, and 458 siMYH7 (2) myofibrils. Details of quantification found in Figure 2M-O.

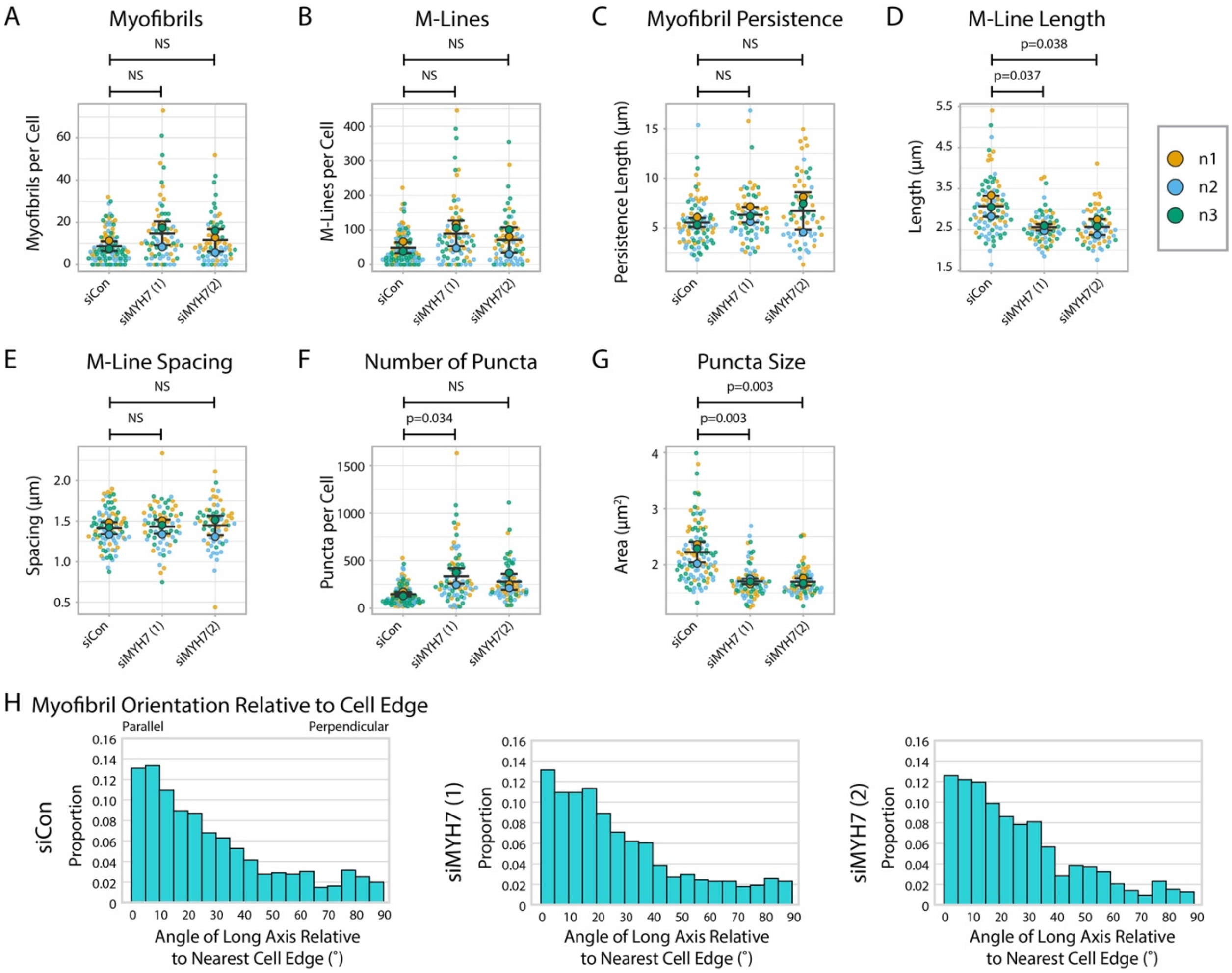


**Figure 7-figure supplement 6: Myomesin quantification and organization in MYH7 knockdown cells**

A) Myofibrils per cell in siCon (scramble)-treated hiCMs and two separate MYH7 siRNA-treated hiCMs (pools 1 and 2). Myofibrils defined as having 3 or more M-Lines in a row. N=3 biological replicates, 104 siCon cells, 75 siMYH7 (1) cells, and 70 siMYH7 (2) cells. Quantification scheme details in Figure 2-figure supplement 1. B) M-Lines per cell in hiCMs from (A). C) Average myofibril persistence length per cell in hiCMs from (A): N=3 biological replicates, 87 siCon cells, 65 siMYH7 (1) cells, and 62 siMYH7 (2) cells. (Only cells with myofibrils were quantified for Figure S13C-E). D) Average M-Line length per cell in hiCMs from (C). E) Average spacing between M-Lines per cell in hiCMs from (C). F) Number of myomesin-positive puncta per cell in hiCMs from (A). G) Average size of all myomesin-positive puncta per cell in hiCMs from (A). H) Myofibril orientation relative to the cell edge segment closest to the myofibril center, perpendicularly. N=3 biological replicates, 793 siCon myofibrils, 775 siMYH7 (1) myofibrils, and 778 siMYH7 (2) myofibrils. Details of quantification found in Figure 2M-O.

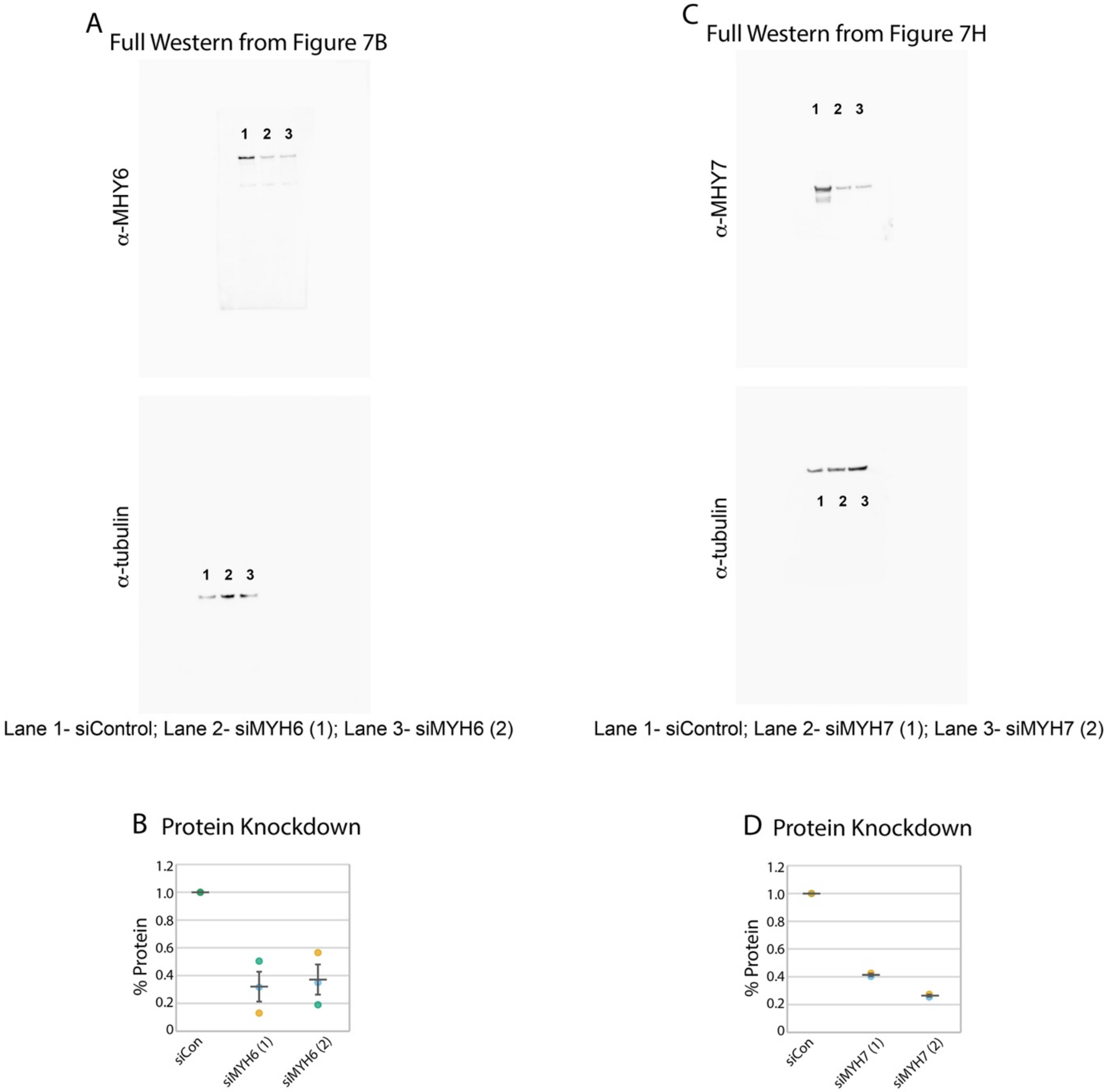


**Figure 7-figure supplement 7: MYH6 and MYH7 knockdown western blots**

A) Full image for the western blots shown in in 7B. Individual lanes are denoted by numbers. Protein knockdown using siRNA targeted to MYH6. C) Full image for the western blots shown in in 7H. Individual lanes are denoted by numbers. D) Protein knockdown using siRNA targeted to MYH7. N=2 biological replicates. Anti-MYH6 and Anti-MYH7 images displayed at 0-50000 gray levels. Anti-tubulin images displayed at 0-40000. Full 16 bit images can be found in Figure 7-source data 1-4.

Figure 7-source data 1: Original 16 bit image file for anti-MYH6 Western blot

See Figure 7-figure supplement 7A

Figure 7-source data 2: Original 16 bit image file for anti-tubulin Western blot

See Figure 7-figure supplement 7A

Figure 7-source data 3: Original 16 bit image file for anti-MYH7 Western blot

See Figure 7-figure supplement 7B

Figure 7-soucrce data 4: Original 16 bit image file for anti-tubulin Western blot

See Figure 7-figure supplement 7B

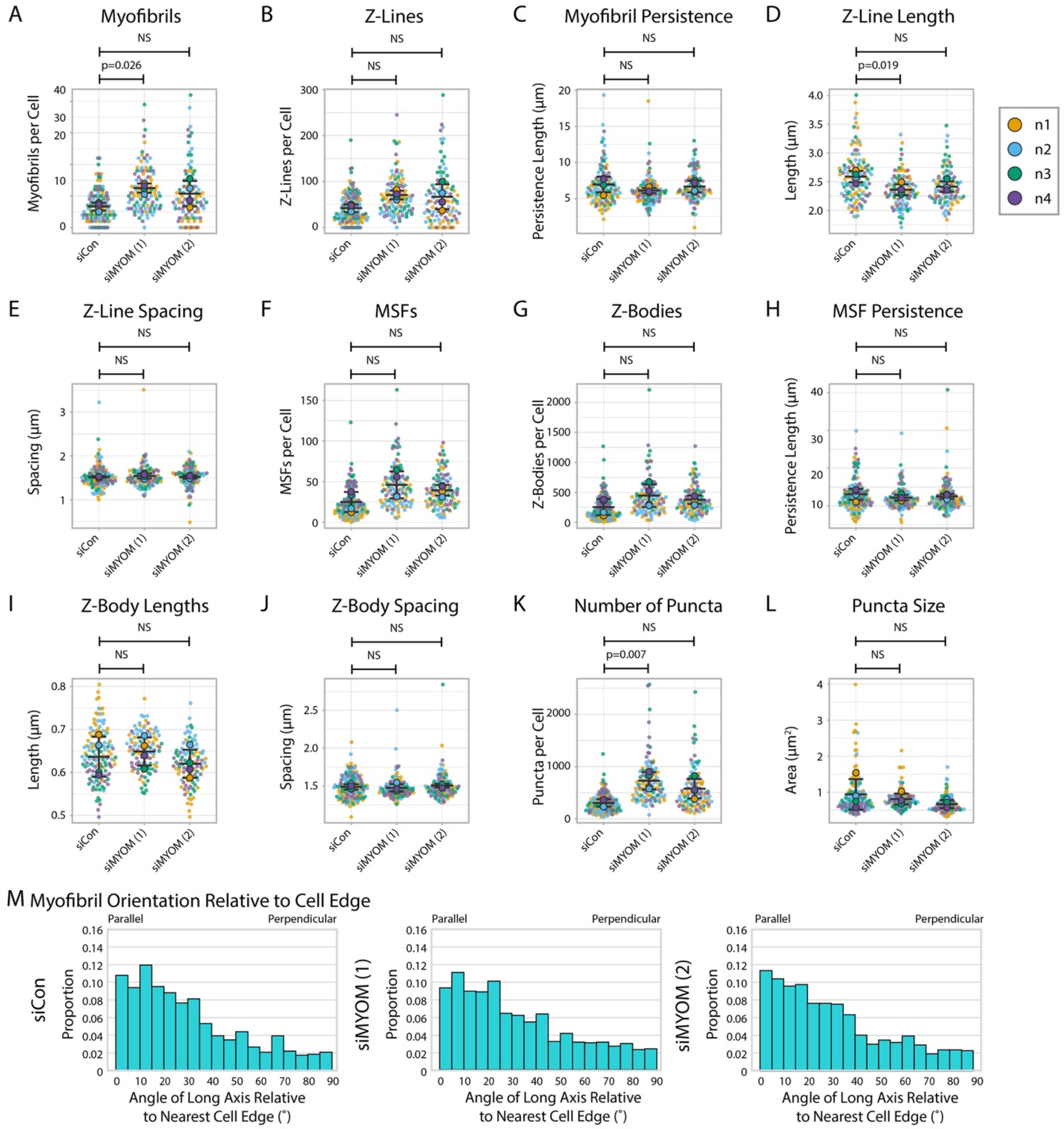


**Figure 8-figure supplement 1: α-actinin-2 quantification and organization in myomesin knockdown hiCMs**

A) Myofibrils per cell in siCon (scramble)-treated hiCMs and two separate myomesin siRNA-treated hiCMs (sequences 1 and 2). Myofibrils defined as having 4 or more Z-Lines in a row. N=4 biological replicates, 132 siCon cells, 105 siMYOM (1) cells, and 103 siMYOM (2) cells. More quantification details found in Figure 2-figure supplement 1. B) Z-Lines per cell in hiCMs from (A). C) Average myofibril persistence length per cell in hiCMs from (A): N=4 biological replicates, 117 siCon cells, 104 siMYOM (1) cells, and 92 siMYOM (2) cells. (Only cells with myofibrils were quantified for C-E). D) Average Z-Line length per cell in hiCMs from (C). E)

Average spacing between Z-Lines per cell in hiCMs from (C). F) MSFs per cell in cells from (A). G) Z-Bodies per cell in hiCMs from (A). H) Average MSF persistence length per cell in hiCMs from Figure S14A: N=4 biological replicates, 131 siCon cells, 105 siMYOM (1) cells, and 103 siMYOM (2) cells. (Only cells with MSFs were quantified for H-J). I) Average Z-Body length per cell in hiCMs from (H). J) Average spacing between Z-Bodies per cell in hiCMs from (H). K) Number of α-actinin-2-positive puncta per cell in hiCMs from (A). O) Average size of α-actinin-2-positive puncta per cell in hiCMs from (A). L) Myofibril orientation relative to the cell edge segment closest to the myofibril center, perpendicularly. N=4 biological replicates, 862 siCon myofibrils, 1305 siMYOM (1) myofibrils, and 1086 siMYOM (2) myofibrils. Details of quantification found in Figure 2M-O.

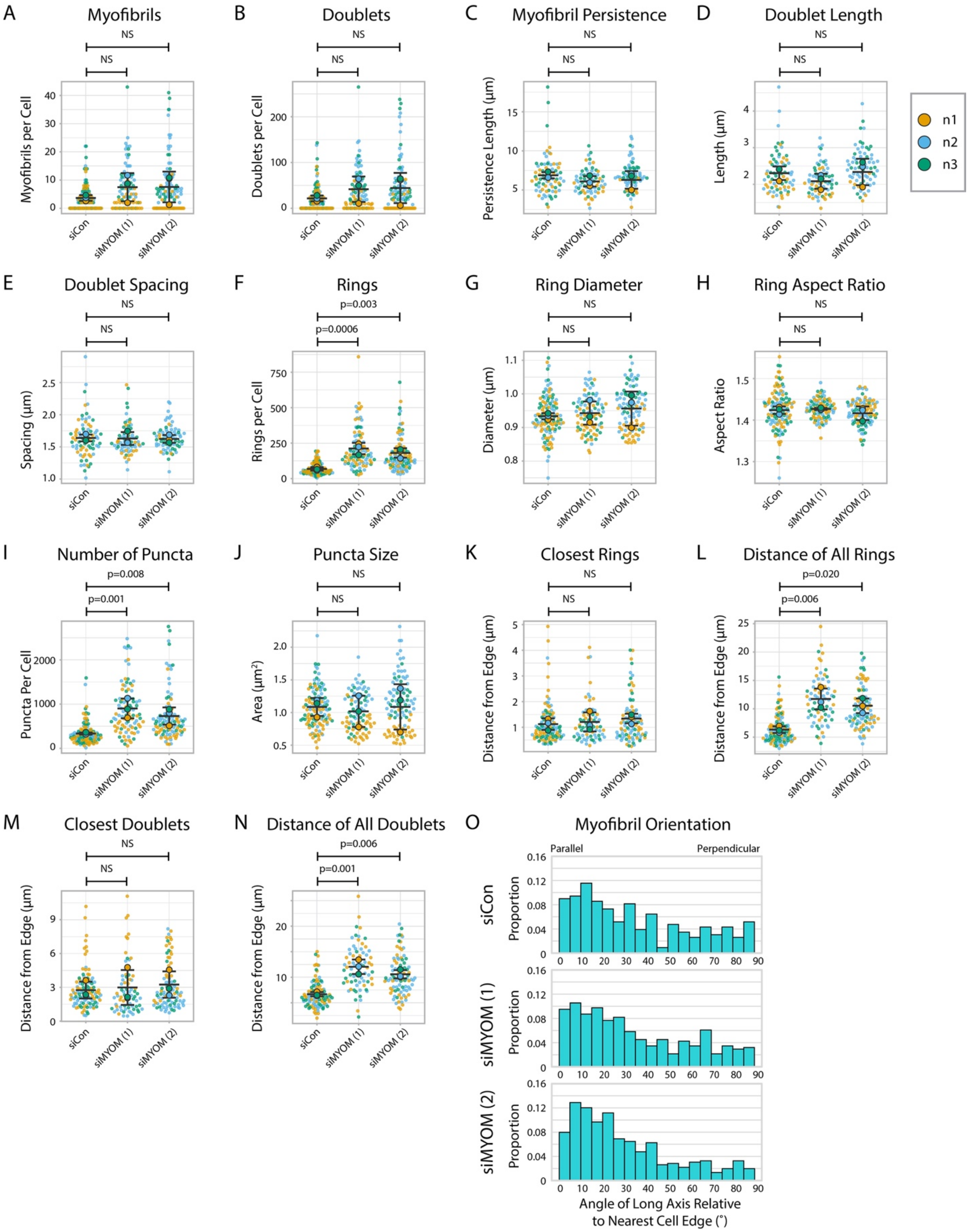


**Figure 8-figure supplement 2: Titin quantification and organization in myomesin knockdown hiCMs**

A) Myofibrils per cell in siCon (scramble)-treated hiCMs and two separate MYOM siRNA-treated hiCMs (sequence 1 and 2). Myofibrils defined as having 4 or more doublets in a row. N=3 biological replicates, 117 siCon cells, 90 siMYOM (1) cells, and 100 siMYOM (2) cells. More quantification details found in Figure 2-figure supplement 1. B) Titin doublets per cell in hiCMs from (A). C) Myofibril persistence length average per cell in hiCMs from (A): N=3 biological replicates, 74 siCon cells, 68 siMYOM (1) cells, and 70 siMYOM (2) cells. (Only cells with myofibrils were quantified for C-E and M-N). D) Average doublet length per cell in hiCMs from (C). E) Average spacing between doublets per cell in hiCMs from (C). F) Number of titin precursor rings per cell in hiCMs from (A). G) Average ring diameter per cell in hiCMs from (A): N=3 biological replicates, 106 siCon cells, 71 siMYOM (1) cells, and 89 siMYOM (2) cells. (Only cells with rings were quantified for G-H and K, L). H) Average ring aspect ratio per cell in hiCMs from (G). An aspect ratio of 1 is circular, and higher ratios are elongated. I) Total number of titin-positive puncta per cell in hiCMs from (A). J) Average size of all titin-positive puncta per cell in hiCMs from (A). K) Average distance from the edge of the five closest titin rings, per cell, in hiCMs from (G). L) Average distance from the edge of all titin rings per cell in hiCMs from (G). M) Average distance from the edge of the five closest titin doublets, per cell, in hiCMs from (B). N) Average distance from the edge of all titin doublets within myofibrils per cell in hiCMs from (B). O) Myofibril orientation relative to the cell edge segment closest to the myofibril center, perpendicularly in hiCMs from (B). N=3 biological replicates, 361 siCon myofibrils, 561 siMYOM (1) myofibrils, and 667 siMYOM (2) myofibrils. Details of quantification found in Figure 2M-O.

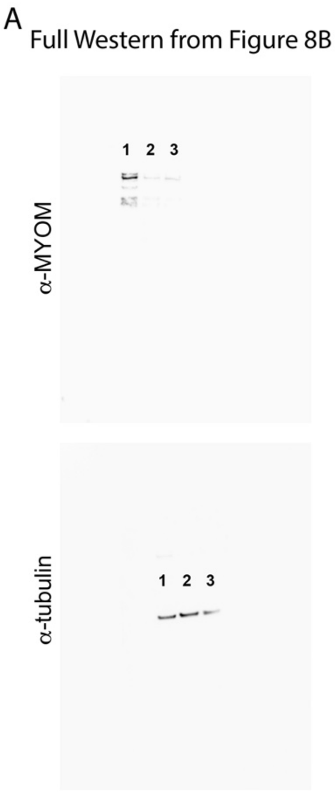


Lane 1- siControl; Lane 2- siMYOM (1); Lane 3- siMYOM (2)

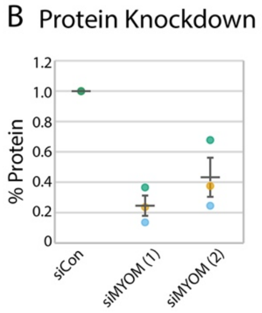


Figure 8-figure supplement 3: MYOM knockdown western blots

A) Full image for the western blots shown in in 8B. Individual lanes are denoted by numbers. Protein knockdown using siRNA targeted to MYOM. C) Full image for the western blots shown in in . Individual lanes are denoted by numbers. D) Protein knockdown using siRNA targeted to MYH7. N=3 biological replicates. Anti-MYOM image displayed at 0-50000 gray levels. Anti-tubulin image displayed at 0-40000. Full 16 bit images can be found in Figure 8-source data 1-2.

Figure 8-source data 1: Original 16 bit image file for anti-MYH6 Western blot

See Figure 8-figure supplement 3A

Figure 8-source data 2: Original 16 bit image file for anti-tubulin Western blot

See Figure 8-figure supplement 3A
