## Supplementary figures and images for "Independent regulation of Z-lines and M-lines during sarcomere assembly in cardiac myocytes revealed by the automatic image analysis software sarcApp"

### Figure 7-source data 1

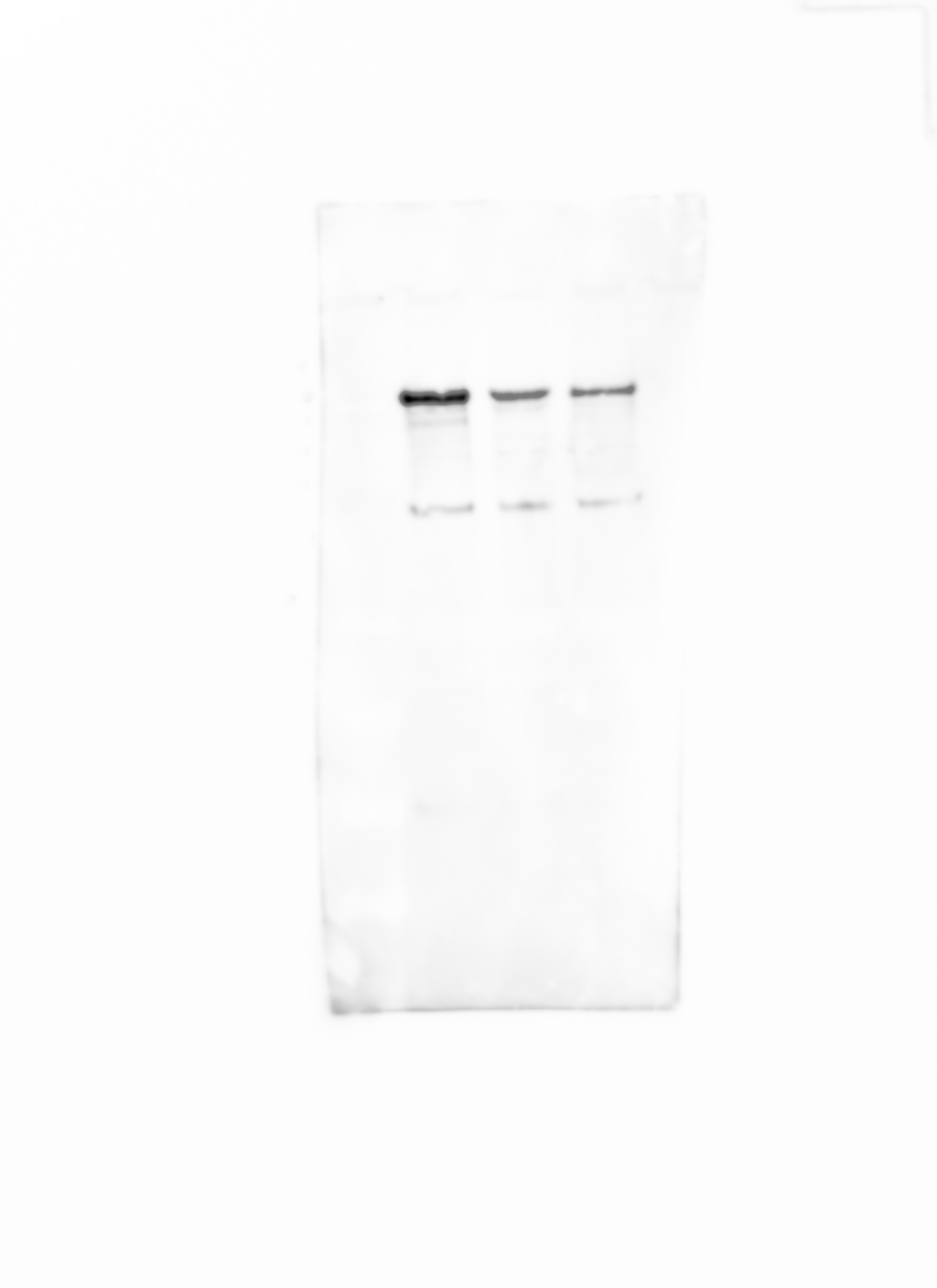

### Figure 7-source data 2

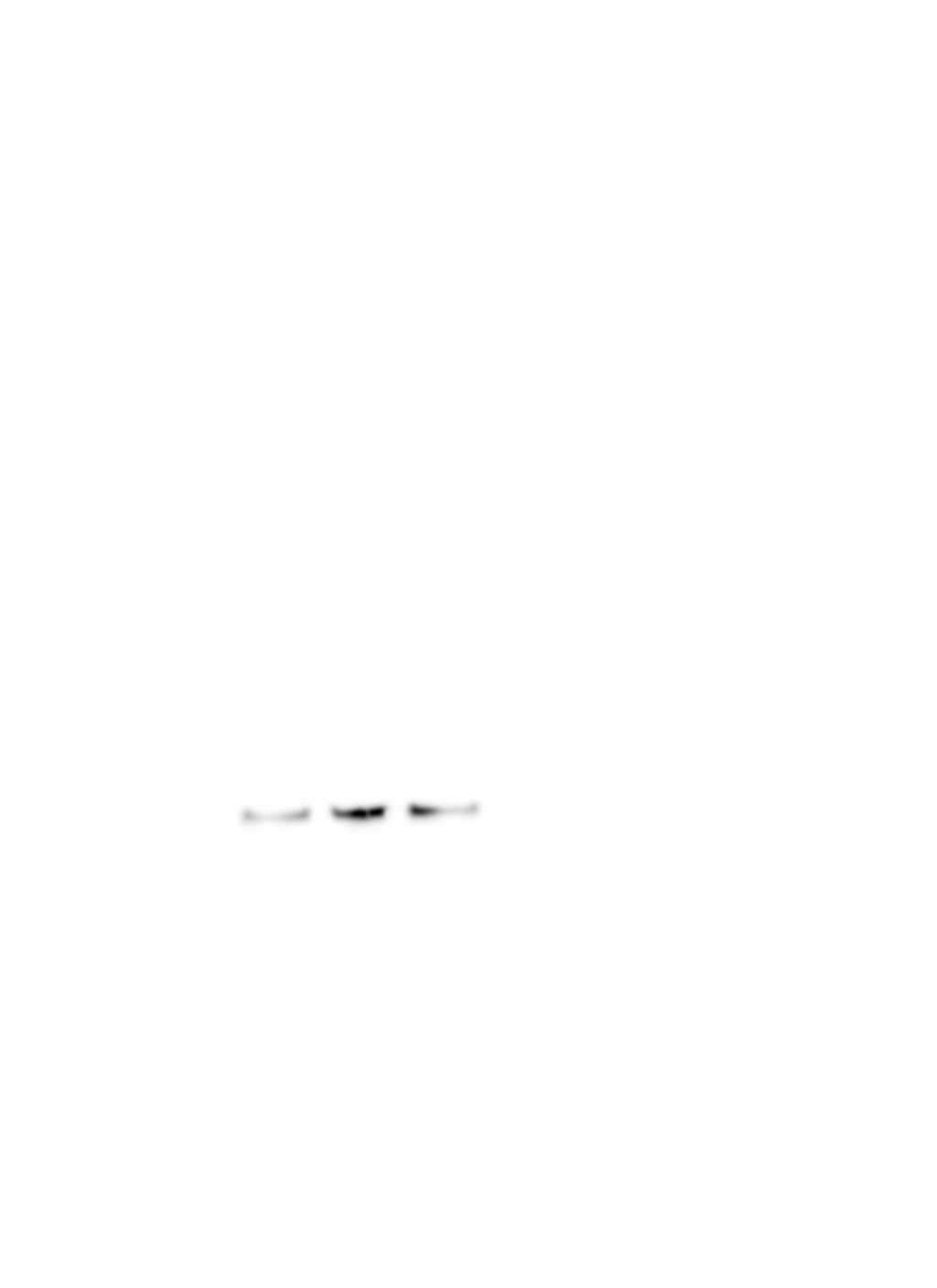

### Figure 7-source data 3

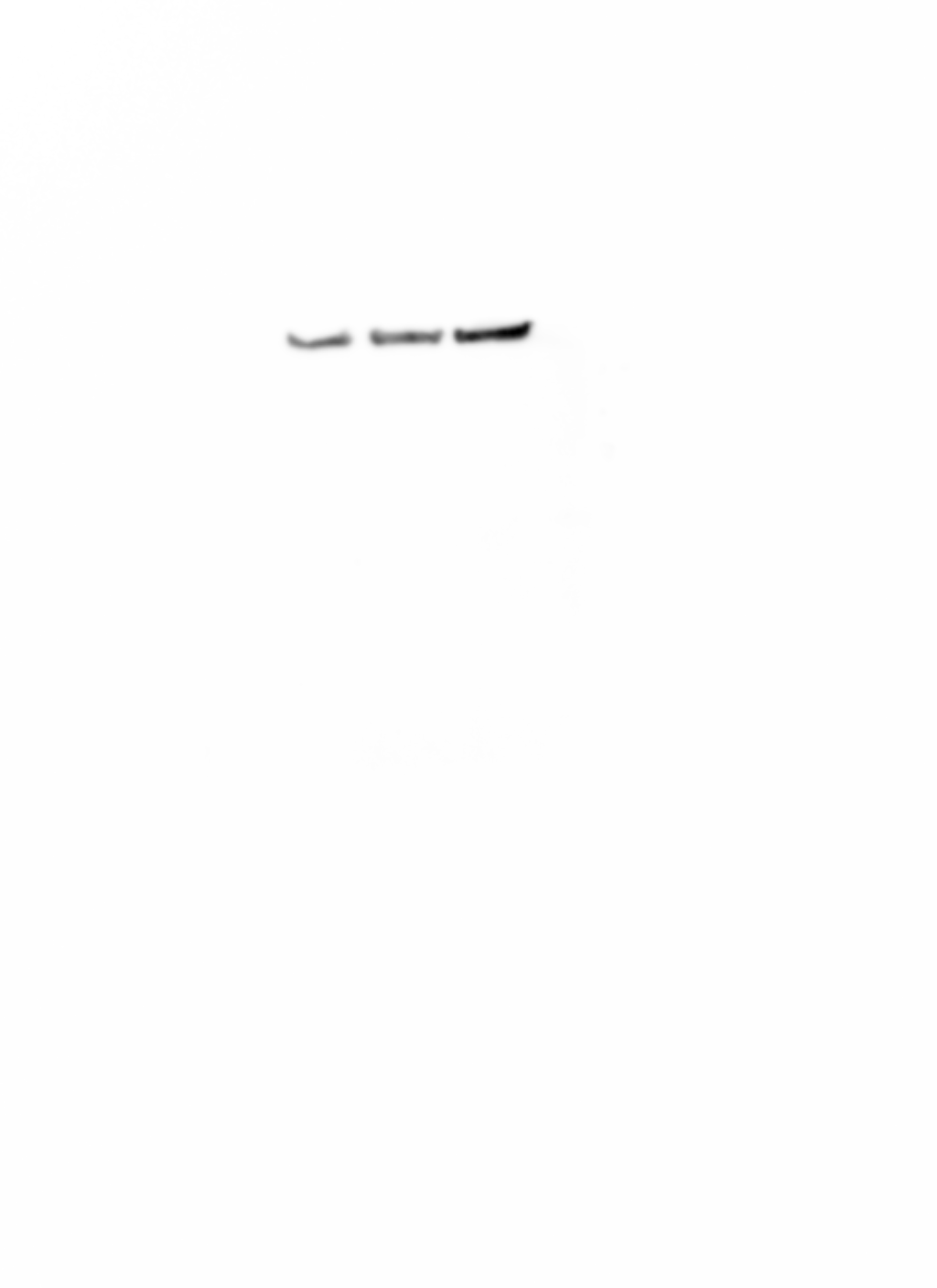

### Figure 7-source data 4

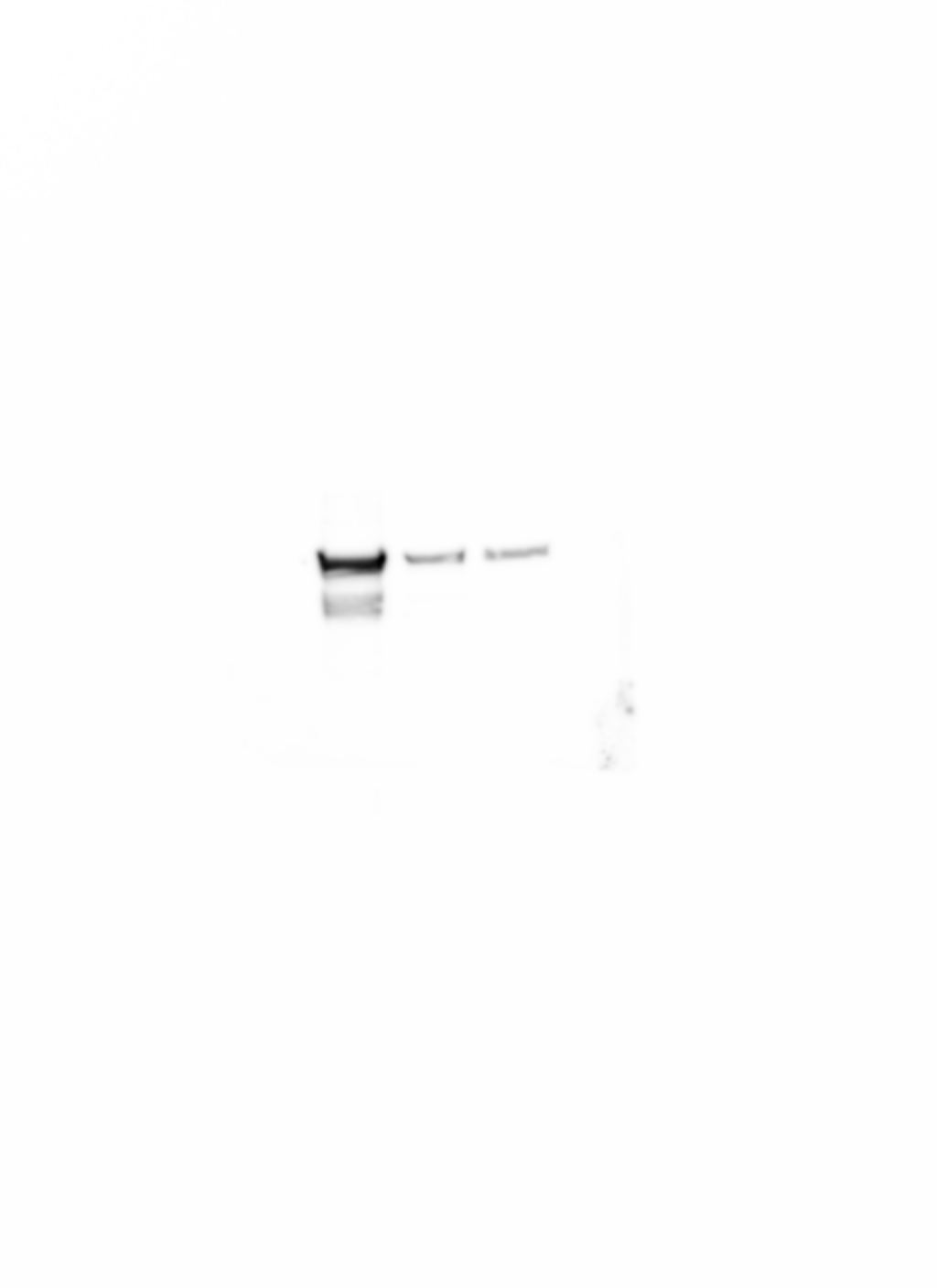

### Figure 8-source data 1

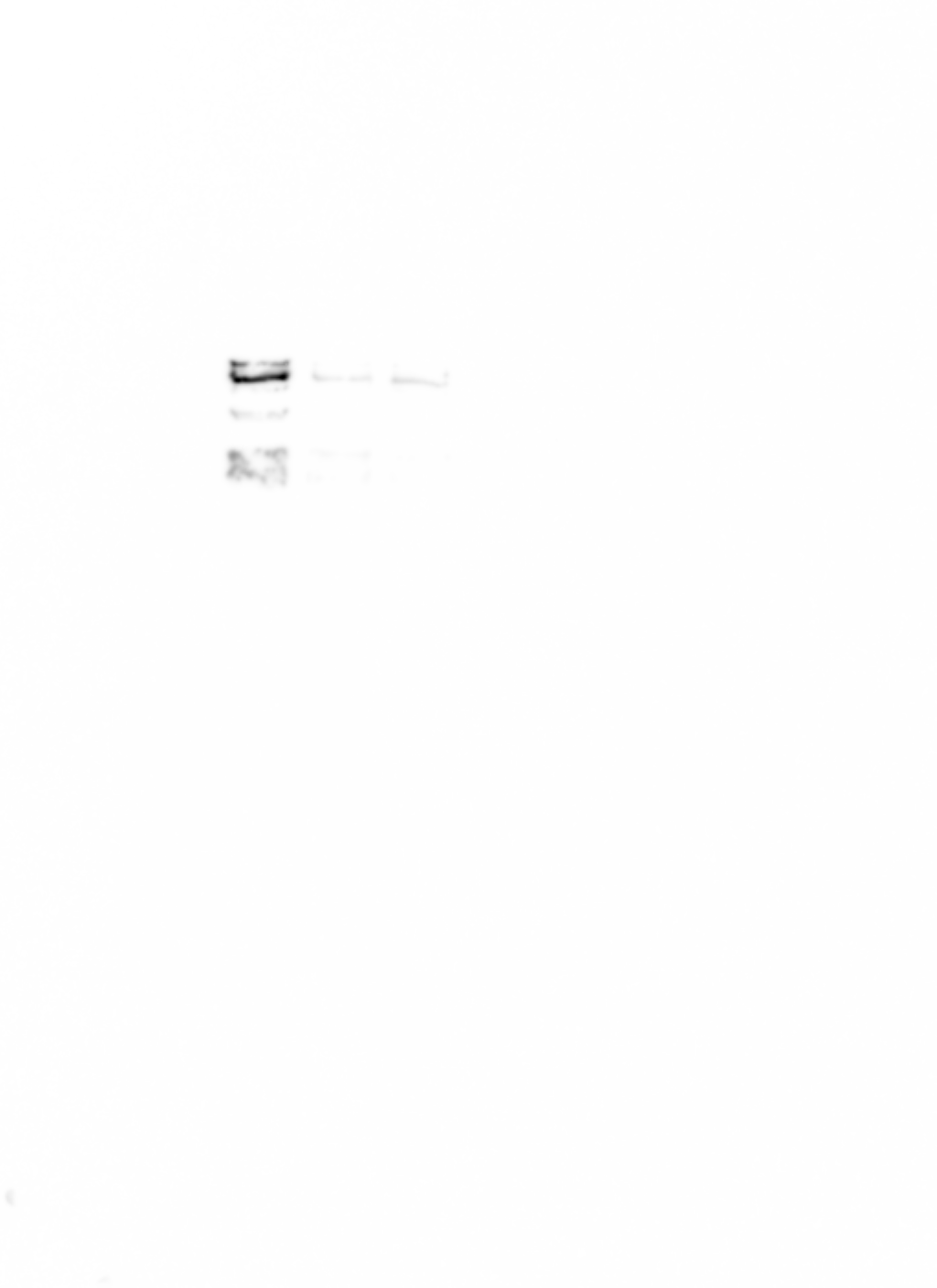

### Figure 8-source data 2

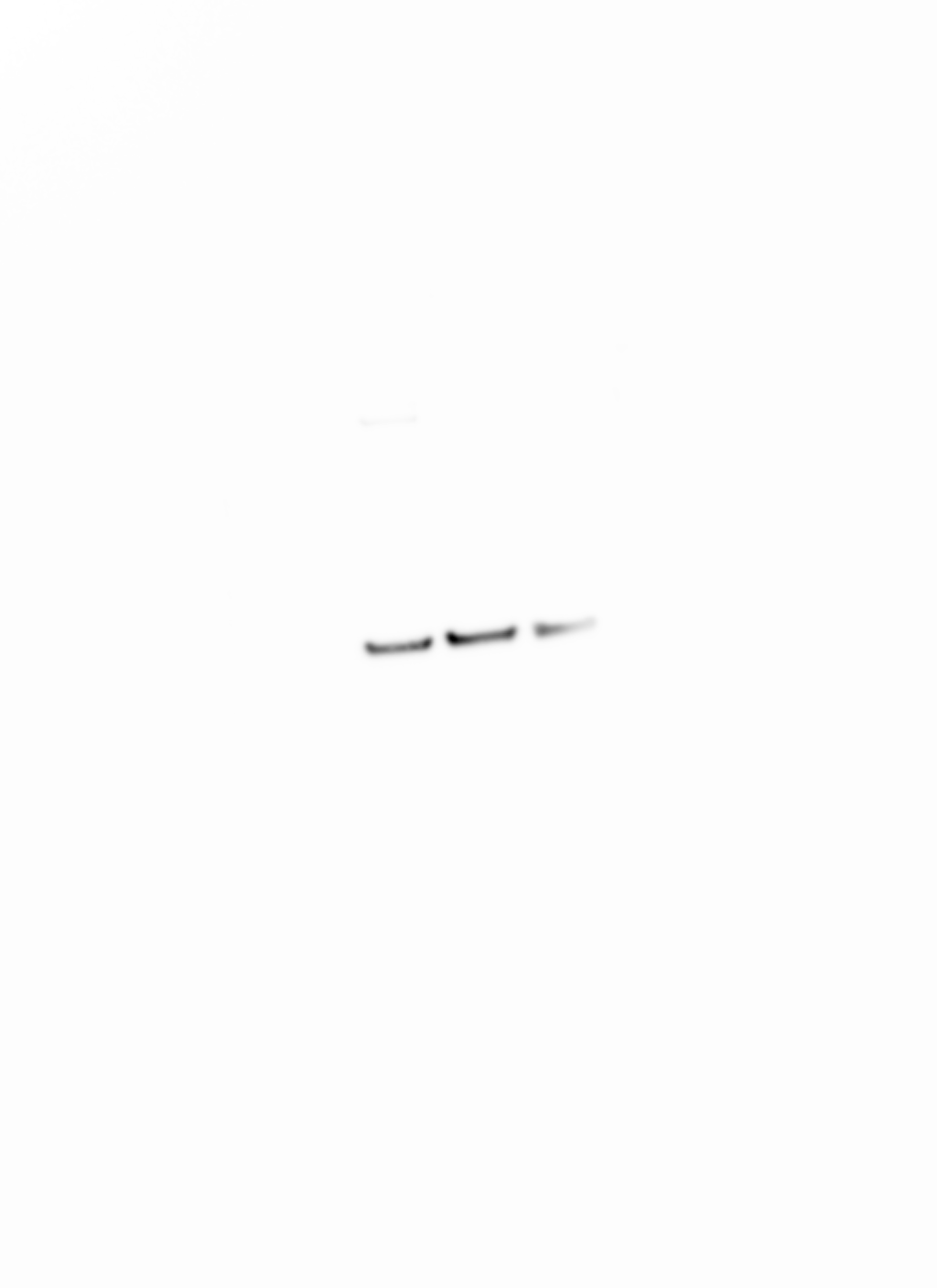
